# Rapid assessment of changes in phage bioactivity using dynamic light scattering

**DOI:** 10.1101/2023.07.02.547396

**Authors:** Tejas Dharmaraj, Michael J. Kratochvil, Julie D. Pourtois, Qingquan Chen, Maryam Hajfathalian, Aviv Hargil, Yung-Hao Lin, Zoe Evans, Agnès Oromí-Bosch, Joel D. Berry, Robert McBride, Naomi L. Haddock, Derek R. Holman, Jonas D. van Belleghem, Tony H. Chang, Jeremy J. Barr, Rob Lavigne, Sarah C. Heilshorn, Francis G. Blankenberg, Paul L. Bollyky

**Affiliations:** Division of Infectious Diseases and Geographic Medicine, Department of Medicine, Stanford University School of Medicine, Stanford, CA 94305, USA; Sarafan ChEM-H, Stanford University, Stanford, CA 94305, USA; Department of Materials Science and Engineering, Stanford University, Stanford, CA 94305; Hopkins Marine Station, Department of Biology, Stanford University, Pacific Grove, CA 93950, USA; Department of Chemical Engineering, Stanford University, Stanford, CA 94305, USA; Felix Biotechnology, South San Francisco, CA, 94080; Division of Gastroenterology and Hepatology, Department of Medicine, Stanford University School of Medicine, Stanford, CA 94305, USA; School of Biological Sciences, Monash University, Clayton, 3800, VIC, Australia; Department of Biosystems, KU Leuven, Leuven, 3001, Belgium; Division of Pediatric Radiology and Nuclear Medicine, Department of Radiology, Lucile Packard Children’s Hospital, Stanford, CA 94305, USA

**Keywords:** bacteriophage therapy, dynamic light scattering, bacteriophage decay, aggregation

## Abstract

Extensive efforts are underway to develop bacteriophages as therapies against antibiotic-resistant bacteria. However, these efforts are confounded by the instability of phage preparations and a lack of suitable tools to assess active phage concentrations over time. Here, we use Dynamic Light Scattering (DLS) to measure changes in phage physical state in response to environmental factors and time, finding that phages tend to decay and form aggregates and that the degree of aggregation can be used to predict phage bioactivity. We then use DLS to optimize phage storage conditions for phages from human clinical trials, predict bioactivity in 50-year-old archival stocks, and evaluate phage samples for use in a phage therapy/wound infection model. We also provide a web-application (Phage-ELF) to facilitate DLS studies of phages. We conclude that DLS provides a rapid, convenient, and non-destructive tool for quality control of phage preparations in academic and commercial settings.

**Significance Statement:** Phages are promising for use in treating antibiotic-resistant infections, but their decay over time in refrigerated storage and higher temperatures has been a difficult barrier to overcome. This is in part because there are no suitable methods to monitor phage activity over time, especially in clinical settings. Here, we show that Dynamic Light Scattering (DLS) can be used to measure the physical state of phage preparations, which provides accurate and precise information on their lytic function – the key parameter underlying clinical efficacy. This study reveals a “structure-function” relationship for lytic phages and establishes DLS as a method to optimize the storage, handling, and clinical use of phages.

## Introduction

As antimicrobial-resistant (AMR) pathogens have become increasingly prevalent (1–3), there is great interest in novel approaches to treating bacterial infections. Bacteriophages (phages), viruses that kill bacteria, offer a promising approach to treating AMR infections that is orthogonal to conventional antibiotics. Phages are safe, well-tolerated (4–10), and potent at sub-nanomolar concentrations (4–6). Phage therapy is already benefiting growing numbers of individual patients (10–12).

However, the development of standardized and reliable phage products—essential for clinical trials and commercial applications—has proved challenging. While long-term storage of phages can be achieved through freezing or lyophilization (13, 14), many phages are often unstable at higher temperatures (e.g. at refrigerated storage at 4 °C or clinical use at 37 °C), which limits their effectiveness (15–20). This has greatly impacted phage therapy clinical trials and commercial development. To this point, poor phage stability was implicated in the case of a recent phage therapy trial for *Pseudomonas aeruginosa* which failed to meet endpoints (9).

Phages, like other multimeric protein assemblies (21, 22), can exhibit poor aqueous stability and decay into non-infectious forms in aqueous solution over time (17). Oxidation contributes to the instability of viral particles (23–26), leading to non-infectious products including aggregates and fragments. For phage therapy and for other research applications involving phages, it is therefore essential to routinely monitor the bioactivity of phages and to optimize conditions for their storage, handling, and transport (13, 14, 20, 27). Moreover, since phages are highly heterogeneous, optimal conditions must be determined separately for each phage using high-throughput techniques, and none currently exist.

Existing methods to assess phage bioactivity/stability (plaque assays) are poorly suited for optimization of storage conditions and routine monitoring of phage preparations. Plaque assays are time-consuming, labor-intensive, destructive to phage stocks, and difficult to scale (18, 28). Rapid, higher-throughput methods to assess phage stability would greatly improve the ability to optimize conditions for phage storage and use.

We hypothesized that Dynamic Light Scattering (DLS), a method for determining the size distribution of nano-scale particles, could yield insights into the physical state of phage particles in solution and thereby serve as an adaptable, high-throughput method to rapidly assess phage stability and bioactivity. We were inspired by previous efforts using DLS to study the impact of ion gradients generated by bacteria on the aggregation/dispersal transitions of T4 phages (29).

Here, we perturb phages with external stressors, including time, temperature, oxidation, ions, and stabilizers. We study phages from human clinical trials and 50-year-old archival phage stocks, finding that external stresses impact the bioactivity of phages and induce aggregation, and that the degree of aggregation as measured by DLS constitutes a quantitative predictor of phage bioactivity, as tested by *in vitro* plaque assays. We also demonstrate how DLS might be used in a workflow to assess the potency of a phage sample for clinical use in a mouse model of wound infection/phage therapy. Building on these insights, we developed a web-based application (Phage-ELF) to enable researchers to predict changes in phage activity from DLS data. These approaches can facilitate screens of phage activity with higher throughput than previously possible.

## Results

### DLS can assess the structural integrity of phages

Our lab and others have observed that phages lose potency in aqueous solution over time at non-freezing temperatures. To explain this phenomenon, we proposed that phages are initially dispersed and functional and over time and in response to environmental conditions, phages fragment into non-infectious particles lacking necessary machinery for infection or aggregate into sterically inhibited clusters (Fig. 1A). Aggregates may also form via denaturation and not contain intact phage particles. We predicted that fragments and aggregates could be observed in small-volume samples of phage preparations (40 μL) and would be distinguishable from intact phages using a standard benchtop Dynamic Light Scattering (DLS) device (Fig. 1B). For these studies, phages were purified from bacterial lysates using standard polyethylene glycol (PEG)-based protocols well-established in the lab, as detailed in the Methods section (30).

**Figure 1.**
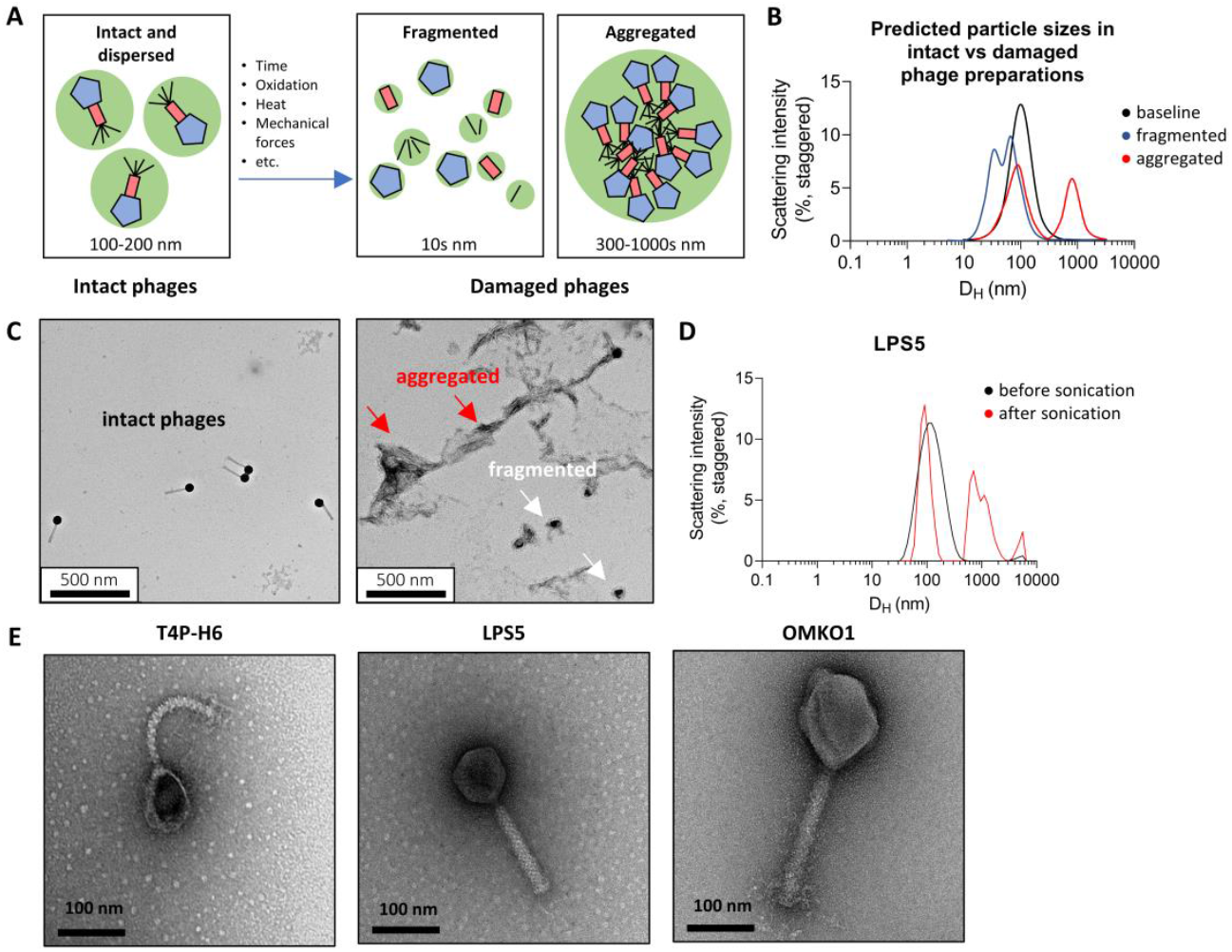
A model of entropic phage decay and assessment by Dynamic Light Scattering (DLS) in phage preparations. **(A)** A model describing mechanisms underlying loss of phage bioactivity. Fragmented and aggregated phage are expected to be distinguishable from intact phage by size. **(B)**, Simulated Dynamic Light Scattering (DLS) spectra representing predicted particle sizes in intact vs. damaged phage preparations. DLS is a benchtop device with a simple workflow that can be used to study the distribution of phages and their byproducts in phage preparations. **(C)** Transmission electron micrographs of intact and sonicated LPS5. The red arrows point to aggregates and the white arrows point to fragments. Shown are representative images obtained at 20,000x magnification. **(D)** Transmission electron micrographs of CYPHY phages, including LPS5. Shown are representative images at 120,000x magnification.

We first measured the DLS spectrum of a phage at baseline and after exposure to conditions expected to cause fragmentation or aggregation. For these studies, we used phage LPS5. LPS5 is one of three antipseudomonal phages used in the Cystic Fibrosis Bacteriophage Study at Yale (CYPHY, NCT04684641; (31–34)). Transmission electron micrographs of CYPHY phages are shown in Fig. 1E, and the physical characteristics of the phages are listed in Table S1.

LPS5 was prepared at a concentration of 10^10^ Plaque Forming Units (PFU)/mL and sonicated for 10 seconds. We analyzed LPS5 before and after sonication by transmission electron microscopy (TEM) and by DLS. As predicted, LPS5 particles were intact and dispersed before sonication. After sonication, phages denatured, with small fragments and large aggregates visible on TEM (Fig. 1C). A similar assessment of phage physical state could be concluded from the DLS spectra. Before sonication, LPS5 produced a single Gaussian peak suggesting a uniform and dispersed population of phages. The hydrodynamic diameter (D_H_) of the phage peak was approximately 100 nm, similar to the actual dimensions of the phage (Table S1). After sonication, the phage peak was lost, and multiple new peaks appeared in the spectra corresponding to fragments with D_H_ = 90 nm, and aggregates with D_H_ = 500-8000 nm (Fig. 1D). These data showed that DLS could be used to measure the physical state of phages, and that phages are intact and dispersed at baseline, and can form aggregates and fragments in response to environmental stress.

### DLS enables monitoring of phage stability over time

Of course, harsh physical manipulations of phages would be expected to cause changes in the size of phage products in solution, and it remained unclear whether observed phage decay over time was also linked to phage aggregation. Therefore, we next applied DLS to investigate the behavior of phages in aqueous suspension over time. We propagated all three CYPHY phages (T4P-H6, LPS5, and OMKO1) and stored them at 10^9^ PFU/mL in conditions in which we generally stored the phages and knew them to be stable (at 4 °C in SM buffer) and in conditions where we expected the phages to be unstable (at 37 °C in PBS). We measured the bioactivity of the phages (reported as phage titer) in terms of plaque-forming units (PFU)/mL and also measured the DLS spectra at baseline and then weekly for 3 weeks.

At 4 °C, all of the phages were stable without loss of titer (Fig. 2A). However, at 37 °C, phages T4P-H6 and LPS5 lost potency – 0.5 logs and 5 logs respectively over the three-week period (Fig. 2B). Correspondingly, the DLS spectra of all of the phages at 4 °C showed no changes (Fig. 2C), and the DLS spectra of T4P-H6 and LPS5 at 37 °C suggested aggregation, with right-shifting of the phage peak and the emergence of large species over time (Fig. 2D). The degree of aggregation appeared to be correlated with the loss of titer, most obviously in the case of LPS5. To quantify changes in the size distribution, we developed a scalar metric we called area-under-the-curve change, or AUCΔ, defined as the non-overlapping area between two DLS spectra. Each DLS spectrum has area of 100, so AUCΔ ranges from 0, representing completely overlapping (identical) spectra, to 200, representing completely distinct (entirely non-overlapping) spectra (Fig. 2E). We measured AUCΔ for all phages over time against the average baseline measurement and found that AUCΔ diverged sharply from baseline under conditions where the phages lost potency (Fig. 2F).

**Figure 2.**
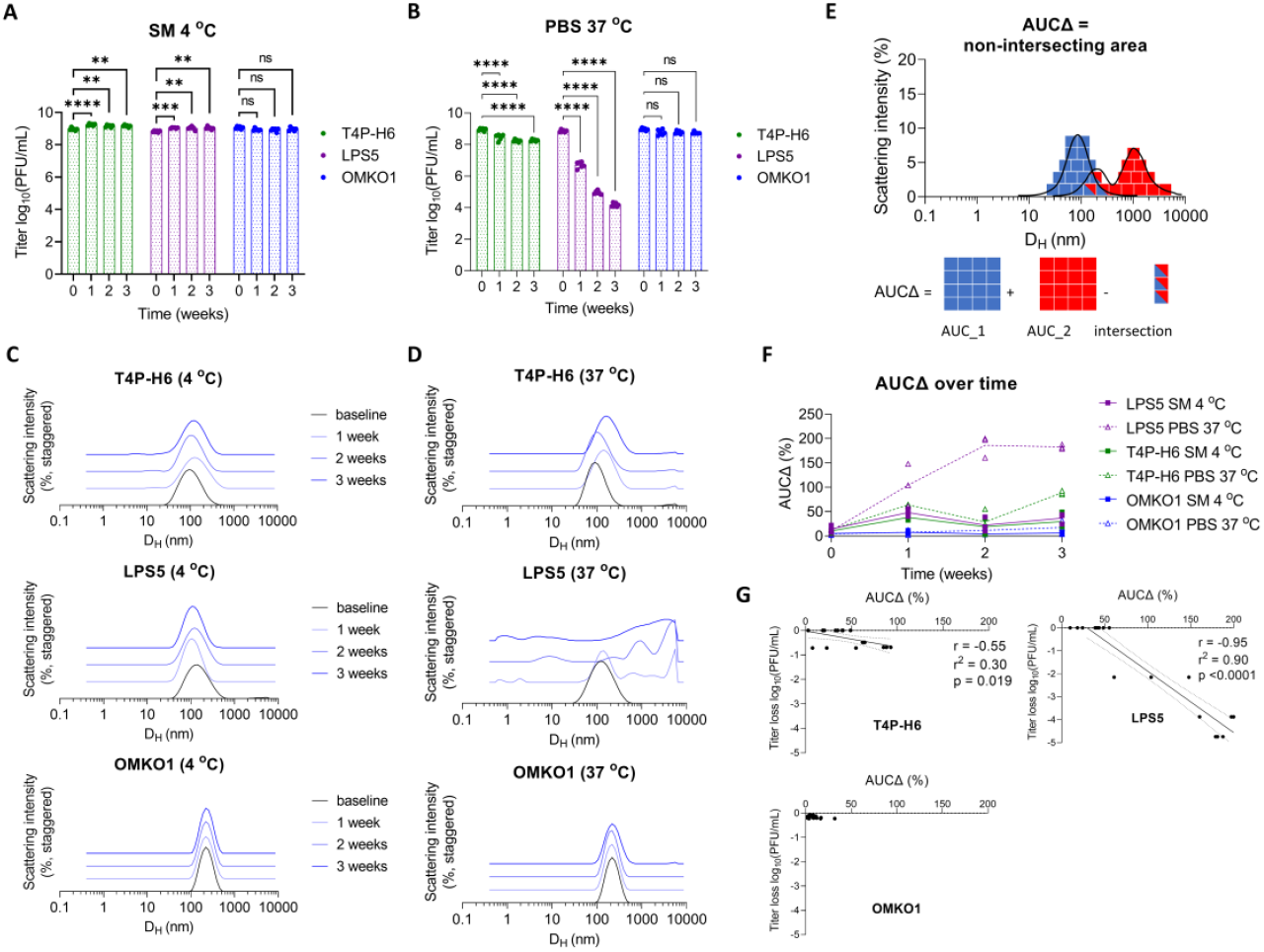
Phages spontaneously aggregate and lose bioactivity in solution over time. Phage preparations can degrade over time in ways that vary between phages. To assess this using DLS, we used a set of phages (T4P-H6, LPS5, and OMKO1) from an ongoing human clinical trial (the CYPHY trial). CYPHY phages (T4P-H6, LPS5, and OMKO1) were stored in stable conditions (SM at 4 °C) or in conditions previously observed to result in titer loss (PBS at 37 °C) for 3 weeks with weekly monitoring by plaque assay and dynamic light scattering (DLS). Titer of CYPHY phages over time in (**A**) SM at 4 °C and (**B**) PBS at 37 °C. Results are from one experiment with n=3 phages, n=6 plaque assays per phage per timepoint. Ordinary two-way ANOVA with Tukey correction, **** = p < 0.0001. (**C, D**) The size-intensity DLS spectra of CYPHY phages over time reveals that aggregation underlies loss of phage activity. Shown are averages of n=3 DLS measurements per timepoint. Darker colors and vertical staggering are used to show progression in time. (**E**) We quantified the non-intersecting area (AUCΔ) between a subsequent DLS measurement and the baseline measurement to quantify changes in size distribution. AUCΔ is a scalar quantity with values ranging from 0 (completely overlapping spectra) to 200 (completely non-overlapping spectra). (**F**) AUCΔ over time for all phages. (**G**) AUCΔ is linearly related to phage titer loss. Two-tailed Pearson’s test for significance of correlation.

AUCΔ and changes in phage titer were linearly correlated, with a strong negative linear correlation between AUCΔ and titer loss for LPS5 in particular (Fig. 2G). 90% of LPS5 titer loss could be explained by changes in the DLS spectrum (i.e., aggregation). These data highlight that changes in the DLS spectrum can indicate loss of phage titer.

We also screened a cohort of 13 phages from Belgium previously used in compassionate use cases. This cohort included phage therapy candidate phages that target *Pseudomonas aeruginosa, Xanthomonas campestris pv. campestris*, and *Staphylococcus aureus* (Table S2). Our previous handling of these phages suggested that some of them were unstable even when stored at 4 °C. We stored these phages at 4 °C in SM buffer and measured their size distributions at baseline and subsequently every three weeks for nine weeks. Many phages showed evidence of aggregation that increased over time (Fig. S1).

Together, these data indicate that an endpoint (and possible mechanism) for loss of phage potency over time in aqueous suspension is aggregation, that aggregation can be tracked using DLS, and that aggregation is closely related to changes in bioactivity. These data underscore that stability measurements need to be made for each phage.

### Optimization of phage storage conditions using DLS

Next, we used DLS to dissect the effects of components of a common phage diluent and storage buffer (SM buffer) on the stability of CYPHY phages. SM buffer is a Tris-buffered solution with NaCl for counterions and osmotic support, often with added MgSO_4_ and gelatin to promote phage stability.

We asked if MgSO4 and gelatin were essential for the stability of CYPHY phages at 37 °C. Therefore, we assessed phage bioactivity and DLS spectrum after 1 week in storage at 37 oC in Tris only, Tris+MgSO4, Tris+gelatin, Tris+MgSO4+gelatin, and Tris+MgSO4+glycerol, compared to 1 week of storage in the same buffers at 4 °C. Glycerol is another common stabilizer for freezing phages, and we wanted to know if it could stabilize CYPHY phages at high temperatures too.

We observed that T4P-H6 was stable in all buffers with no titer loss or change in DLS spectrum (Fig. 3A, D). LPS5 was unstable at 37 °C in buffers lacking MgSO4: Tris only, Tris+MgSO4, and Tris+MgSO4+glycerol, with >2 log titer loss relative to phage stored at 4 °C (Fig 3A). These conditions also showed marked aggregation (Fig. 3E). OMKO1 was unstable at 37 °C in Tris only, losing 1 log (Fig 3C). However, OMKO1 DLS spectra at 37 °C were all similar to 4 °C, suggesting some lack of sensitivity (Fig. 3F).

**Figure 3.**
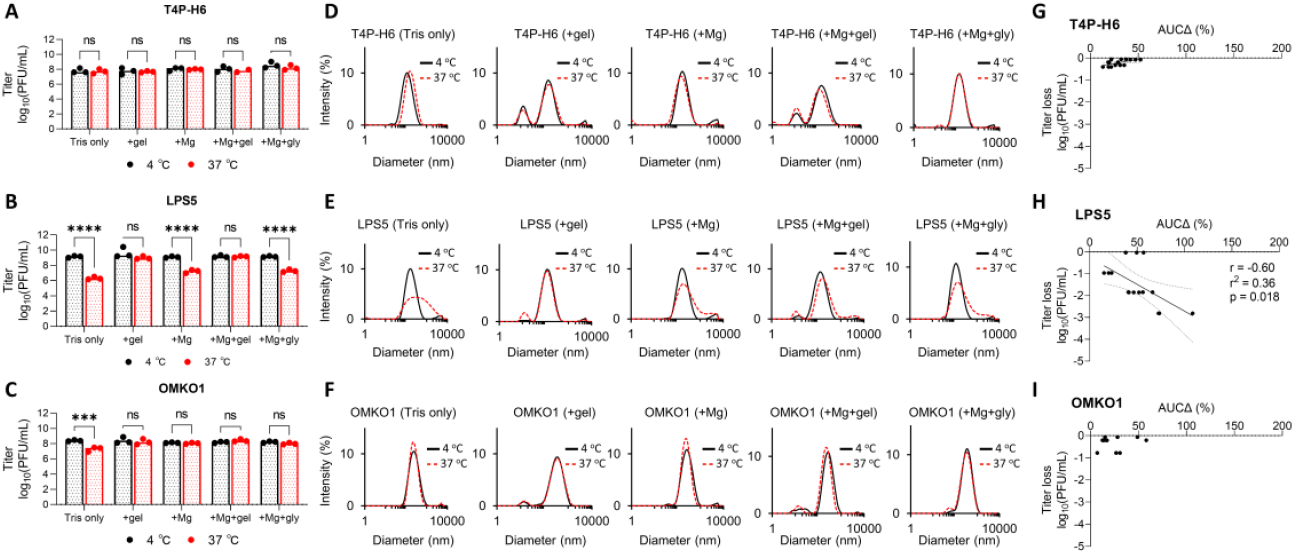
Optimization of phage storage conditions using DLS. Typically, temperature and buffer conditions must be optimized individually for each phage. We assessed this for the CYPHY phages using DLS. (**A** through **C**) Titer of CYPHY phages after 1 week of storage in several buffers (Tris, Tris+gel, Tris+Mg, Tris+Mg+gelatin (SM buffer), and Tris+Mg+glycerol) at 4 °C or 37 °C for 1 week. n=3 plaque assays were performed for three phages for five buffer conditions each. Ordinary two-way ANOVA with Tukey correction. *** = p<0.001, **** = p<0.0001. (**D** through **F**) DLS spectra of CYPHY phages in different buffers. The black line represents phage stored at 4 °C, while the red dashed line represents phage stored at 37 °C. Shown are averages of n=3 DLS measurements for each phage and buffer condition. (**G** through **I**) AUCΔ vs titer loss for CYPHY phages. Data are shown from one experiment. Two-tailed Pearson’s test for significance of correlation.

Temperature and buffer selection are important variables that impact phage stability and function. These data support the use of DLS to select optimal conditions and identify necessary additives to support phage stability under desired conditions.

### Oxidative damage to phages is reflected in DLS measurements

We next investigated the relationship between AUCΔ and titer loss following oxidation. Our rationale for this was two-fold. First, we wanted to simulate oxidative stress that occurs in aqueous suspension over time (23) or in tissues (35). Second, we wanted to test the linear relationship AUCΔ and titer loss in a controlled system. For this experiment, we prepared CYPHY phages at 1.25 x 10^8^ PFU/mL, titrated the phages with 0-20% H_2_O_2_ for 20 minutes at room temperature, and then removed the unreacted H_2_O_2_ by several rounds of dialysis. We recovered the phage from the dialysis cassettes and measured the resultant titer loss by plaque assay and changes in size by DLS and TEM.

We observed that phages were variably sensitive to oxidative stress. T4P-H6 was most resistant to oxidation, with no titer loss with <2% peroxide, and ∼1 and ∼3 log loss with 2% and 20% H_2_O_2_ respectively (Fig. 4A). Correspondingly, aggregates were observed in the DLS spectrum at 2% and 20% H_2_O_2_ (Fig. 4B). LPS5 suffered 1-log, 4-log, and complete titer loss in response to 0.2%, 2%, and 20% H_2_O_2_ (Fig. 4A). Right-shifting of the DLS-spectrum was observed at 0.2% and 2% H_2_O_2_, with marked aggregation at 20% (Fig. 4C). OMKO1 was most sensitive, suffering 1-log titer loss at 0.2% peroxide and complete loss thereafter (Fig. 4A). The DLS spectra were unchanged at 0.2% H_2_O_2_, but aggregates were visualized at higher peroxide concentrations (Fig. 4E). TEM showed that phages were intact and dispersed at baseline, but then formed aggregates upon oxidation with 20% H_2_O_2_ (Fig. 4, E through J).

**Figure 4.**
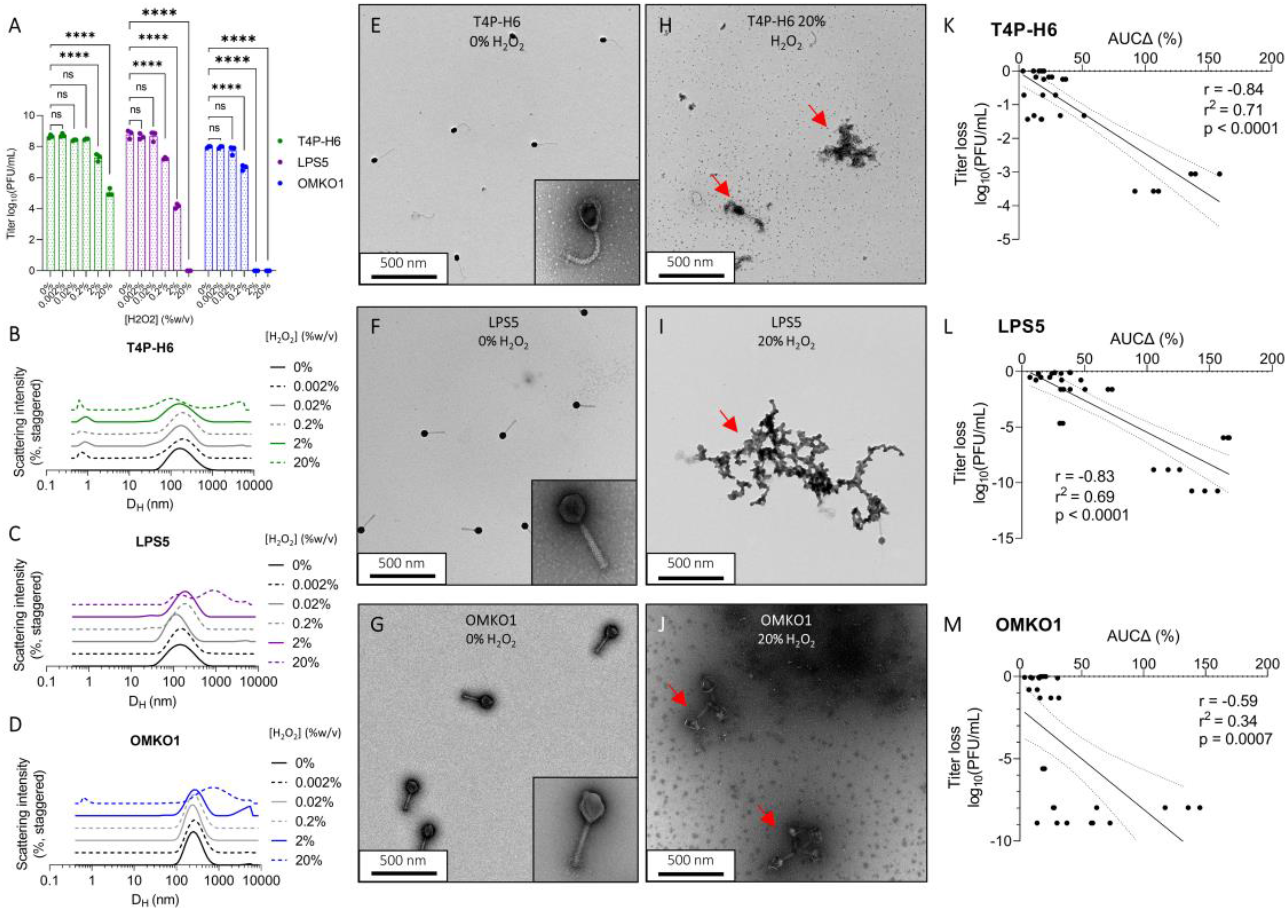
Oxidation drives aggregation of phage, which predicts loss of bioactivity. We assessed the impact of oxidative damage on the CYPHY phages using DLS and functional assessments of phage titer. (**A**) Bioactivity titer of CYPHY phages in response to titration with 0-20% H_2_O_2_. Titer was measured by n=3 plaque assays per concentration. Titration was representative of n=2 independent experiments. Ordinary two-way ANOVA with Tukey correction, **** = p < 0.0001. (**B** through **D**) DLS spectra of titrated phages. Shown are averages of n=3 DLS measurements for each peroxide concentration. Note peroxide concentration-dependent changes in size. (**E** through **J**) Transmission electron micrographs of (**E** through **G)** intact and dispersed CYPHY phages (inset: a single phage at 120,000x magnification) and (**H** through **J**) aggregated CYPHY phages. Shown are representative images obtained at 20,000x magnification. (**K** through **M**) AUCΔ vs titer loss for CYPHY phages a high proportion of the variation in titer is explained by variation in AUCΔ (i.e., aggregation). Linear regression includes data pooled across n=2 independent experiments. Two-tailed Pearson’s test for significance of correlation.

We measured AUCΔ for all phages and was compared to titer loss by simple linear regression. We observed moderate to strong negative correlations between AUCΔ and titer loss for all CYPHY phages (Fig 4, K through M). We additionally observed that variation in AUCΔ explained a high proportion of the variation in titer (r^2^ = 0.71, 0.69, and 0.34 for T4P-H6, LPS5, and OMKO1 respectively).

Together, these data indicate that oxidative damage impacts phage bioactivity. Oxidative damage is reflected in DLS measurements as aggregation, and changes in size distribution are directly related to titer loss.

### DLS measurements reflect changes in lytic function in 50-year-old T-series phages

Next, we used DLS to evaluate the status of phages that had been stored for long periods. For this, we evaluated five “T-series” *Escherichia coli* phages that were stored in sealed glass ampules at 4 °C over 50 years, starting in 1972 (36) (Fig. 5A). Previous work identified that these phages had lost nearly all of their bioactivity, except phage T6 (36) (Fig. 5B).

**Fig. 5.**
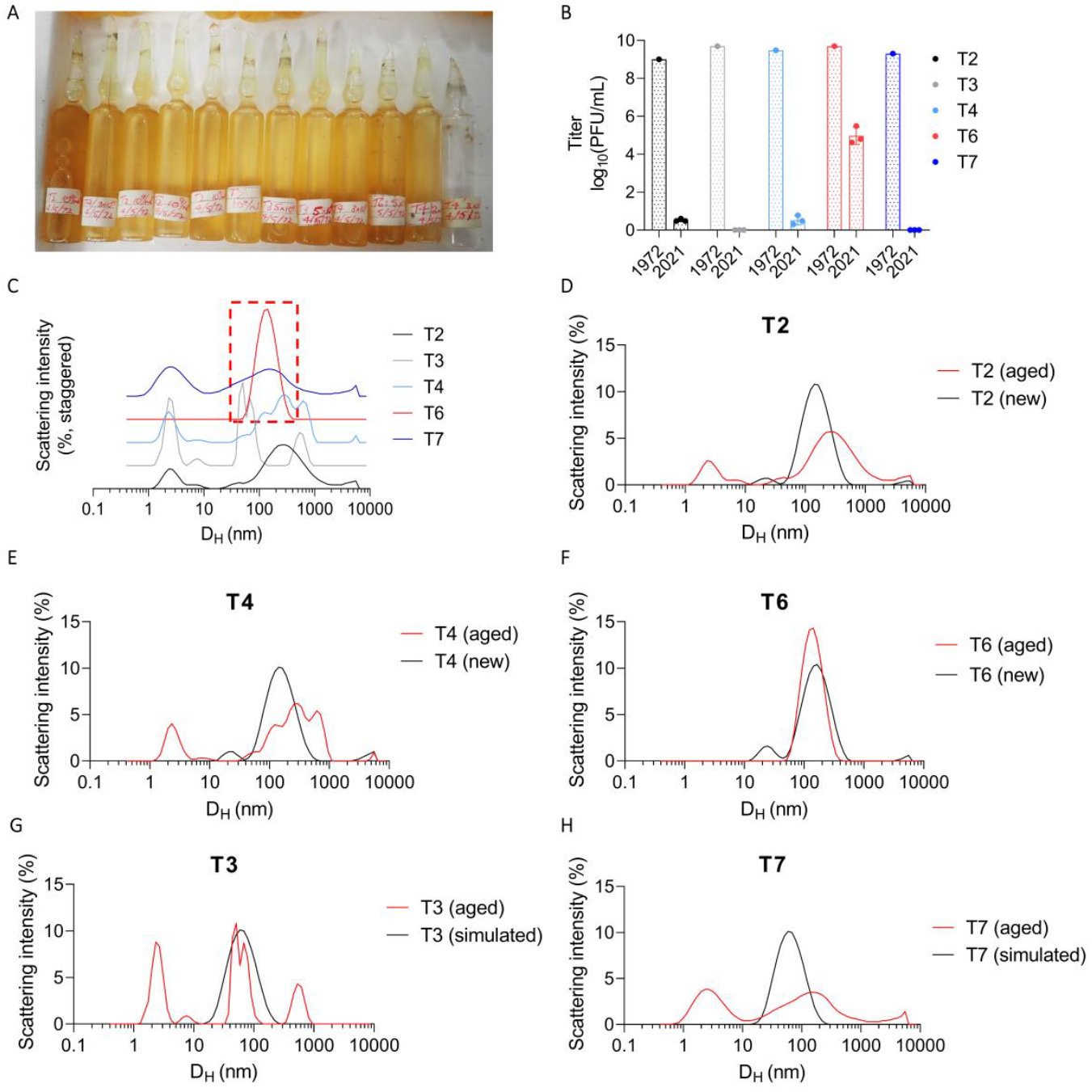
DLS accurately identifies active phage in a cohort of inactive phages stored for 50 years. To evaluate the ability of DLS to assess the stability of phage preparations, we studied a set of archival phage stocks. (**A**) Picture of glass ampules in which T-series phages were stored for five decades. (**B**) Titer of n=5 T-series phages prior to storage in 1972 and after opening in 2021. Here, we include titer data from Subedi and Barr (*36*). 1972 data is reported as a single measurement. 2021 data is reported as n=3 plaque assays per phage. (**C**) DLS spectra of T-series phages measured in 2022. Shown are averages of n=3 DLS measurements per phage. Only T6 remains as a monodisperse Gaussian with a mean hydrodynamic diameter of ∼100 nm and was accordingly the only phage to retain any activity. (**D** through **H**) DLS spectra of old vs. newly propagated phage or simulated DLS spectra. Shown are averages of n=3 DLS measurements per phage.

We used DLS to obtain the size distribution of the T-series phages. Most phage samples were highly polydisperse, reflecting both aggregates and fragments. The exception was phage T6, which produced a single Gaussian peak at 200 nm (Fig. 5C). We propagated fresh samples of T2, T4, and T6 to perform a comparison between an aged sample and a fresh one. This revealed substantial differences in the spectra in the case of T2 and T4, but not in the case of T6 (Fig. 5, D through F). We were unable to propagate T3 and T7, so we predicted the baseline spectra for these phages as Gaussians centered on D_H_ = 60 nm. This predicted significant differences in their spectra between aged and fresh samples (Fig. 5, G and H).

These data indicate that DLS can be used to distinguish active from inactive phage samples and to identify active phage stocks in a collection.

### Genomic damage is not reflected on DLS measurements

We predicted that environmental conditions that drive genomic damage alone would cause phage titer loss but not changes in phage size distribution and hence would not be reflected on the DLS spectra. To this end, we irradiated CYPHY phages as well as LUZ19 and LUZ24 from the Belgian cohort with germicidal UV-C light (254 nm) for 20 minutes, and measured the resultant titer loss by plaque assay and size distribution by DLS.

We observed that irradiation resulted in near-total loss of activity for all phages (Fig. S2A), but did not affect their size distribution (Fig. S2, B through F). Together with the data in previous sections, these data indicate that structural but not genomic damage drives phage aggregation, and that DLS is best used to assess phage titer loss in samples that have been shielded from ionizing radiation.

### DLS can prospectively evaluate phage samples for clinical use and inform *in vivo* phage therapy

We next modeled how DLS can be used to evaluate phage preparations for clinical use. We anticipate that prospective evaluation of phage samples could serve as a useful quality-control step before administration to patients. To this end, we first developed a web-based software application called “Phage-Estimator of Lytic Function (Phage-ELF; URL: https://jp22.shinyapps.io/shinyapp/). This tool accepts minimally processed DLS and lytic activity data in the.csv format to visualize DLS curves, calculate AUCΔ, train a linear or logistic model, and finally make predictions on new DLS data (Fig. 6A).

**Figure 6.**
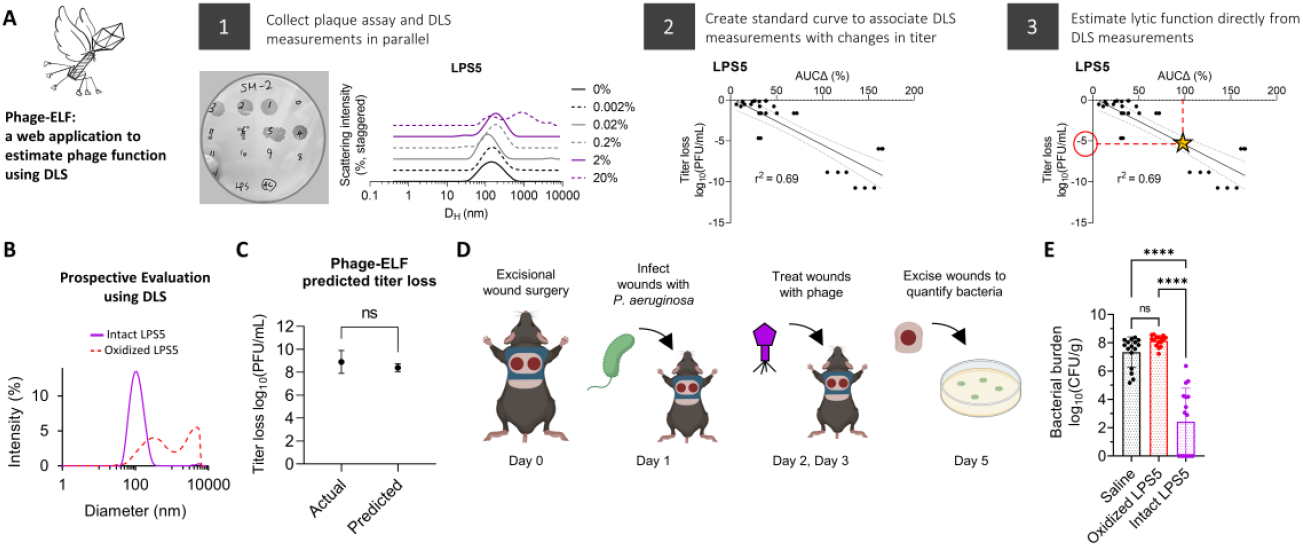
DLS-based predictions of lytic activity can inform the success of *in vivo* phage therapy. We developed a web-based application (Phage-<u>E</u>stimator of <u>L</u>ytic <u>F</u>unction (Phage-ELF)) to assess phage titer loss using a standard curve generated for a given phage strain and set of perturbations. (**A**) Phage-ELF trains a linear model on paired DLS and lytic activity data, which can then be used to predict titer loss based on new DLS data and our ΔAUC analysis. This application and sample data are available for download directly from https://jp22.shinyapps.io/shinyapp/. (**B**) DLS spectra of intact and oxidized LPS5. (**C**) Phage bioactivity titers predicted from DLS data and Phage-ELF and actual measurements from lytic plaque assays. Two-tailed two-sample Student’s t-test. (**D**) Schematic of topical phage therapy in mouse model of *Pseudomonas aeruginosa* wound infection. Mice were treated with saline, intact LPS5 (10^10^ PFU/mL), or oxidized LPS5 on day 1 and 2. Shown are averages of n=3 DLS measurements per phage. (**E**) Bacterial burden after treatment. One-way ANOVA with Tukey correction, **** = p < 0.0001. Results from n=16 wounds for n=3 treatment conditions. Data are representative of n=2 independent experiments.

We prepared paired samples of LPS5 at 10^10^ PFU/mL and inactivated one by oxidation. The identity of each sample was blinded to the experimenters. Then, DLS spectra of each sample was measured, which identified one sample to have a single Gaussian peak and the other to be severely aggregated (Fig. 6B). Using the calibration curves for oxidized LPS5 phage, Phage-ELF estimated the titer loss to be 8.4 ± 0.3 log_10_PFU/mL. This closely compared to the observed titer loss of 8.9 ± 1.0 log_10_PFU/mL that was later measured (Fig. 6C).

To further assess the functional properties of these samples, we used a mouse model of phage therapy. This incorporated a delayed inoculation model of *P. aeruginosa* infection previously published by our lab (37–39). Briefly, anesthetized mice received full-thickness bilateral dorsal wounds made using a 6 mm biopsy punch. The wound was then covered and inoculated the following day with 10^3^ colony-forming units (CFU) of bacteria. Two hours after infection and after the inoculum had been absorbed into the wound bed, mice were treated with saline, the phage sample predicted to be active, or the phage sample predicted to be inactivated. Three days post infection, bacterial infections were assessed by plating wound homogenates for CFUs (Fig. 6D).

We observed that treatment with the intact phages resulted in bacterial eradication in 50% of wounds and a significant reduction in colonization relative to control and oxidized phages (Fig. 6E). These data demonstrate that DLS predictions of phage titer loss were precise in this model, and that DLS can provide useful, clinically relevant information about the activity of a phage sample prior to its use.

## Discussion

We report that DLS tracking of phage size distributions is an effective way to predict the bioactivity of phage preparations in a semi-quantitative to quantitative manner. We demonstrate that many environmental factors and handling procedures cause phage damage and titer loss, with aggregation being an associated endpoint. These findings were made using phages from human clinical trials and 50-year-old archival phage stocks. Thus, phage bioactivity is impacted in ways that can be predicted from their physical state.

DLS offers several advantages over plaque assays for assessing phage stability. Once standard curves are developed to associate changes in physical state and changes in bioactivity, DLS alone is faster, higher-throughput, and less destructive than plaque assays. Our vision is that this tool will enable optimization of storage conditions for individual phages and quality control for post-production batches of phages intended for clinical and research applications. Because DLs instruments are already common laboratory equipment, the approach developed here could be readily adapted to commercial and research settings where monitoring phage stability is essential.

Similar tools are suggested for product development and process analytics in gene therapy programs involving adenovirus (40). However, these approaches have not made inroads into phage therapy, perhaps because the heterogeneity of individual phage preparations and morphologies, and because of the critical importance of lytic activity as an endpoint. We show here that physical state can serve as an effective proxy of lytic activity, for phages with diverse morphologies.

To encourage and highlight the translational potential of this tool, we developed a web-based application to facilitate DLS analyses of phages, and demonstrated that DLS-based predictions of phage activity can inform *in vivo* phage therapy. For these studies, we established *P. aeruginosa* wound infections in mice using a delayed inoculation protocol and subsequently administered phage therapy. While animal models have previously been used to study phage therapy for wound infections (41), these typically co-administer bacteria and phages or involve indolent organisms or immunocompromised hosts. Our protocol, using conventional C57BL6 mice and a major cause of human wound infections *(P. aeruginosa)*, will enable future work in this field. Our simple algorithms can be used to assess titer change using a standard curve for a given phage and set of perturbations, and we hope that these studies and tools will enable future therapeutic studies and research into the structure-function relationships of lytic phages.

Our approach has several limitations. First, calibration curves linking particle size to lytic activity must be re-derived for each phage and set of conditions. Second, the DLS spectra are sensitive to the sampling procedure employed – the measured sample must contain the aggregates, fragments, and intact components of the population. Third, DLS is insensitive to sources of direct genomic damage such as UV light. Fourth, it is likely that DLS cannot distinguish “ghost” phage particles (i.e., virions that have expelled their genomes) from active phage particles. While phages are typically stored in ways that protect them from UV light; nonetheless, additional assays may be required to specifically control for these issues.

Many questions remain. Our studies identified that phages had different stability profiles, but the mechanisms underlying differential phage stability remain unknown. The relevance of these approaches to phage cocktails awaits further investigation. These studies may also benefit from including more diverse phages and other viruses (e.g. adenovirus).

## Materials and Methods

### Bacteriophage isolation and propagation

Phages and their bacterial hosts used in this study are listed in Table S2. CYPHY phages were a kind gift from Felix Biotechnology. Belgian cohort phages were a kind gift from Rob Lavigne and Jean Paul Pirnay. T-series phages were a kind gift from Jeremy Barr.

Phages were propagated using techniques that are well-established in our lab(38). Briefly, we infected mid-log or early-log phase planktonic bacterial cultures phage until clearing was observed relative to a non-infected culture. We then removed gross bacterial debris by centrifugation (8000 x *g*, 20 min, 4 °C), and filtered the supernatant through a 0.22 μm polyethersulfone (PES) membrane (Corning, Corning, New York, Product #4311188). The supernatant was treated with 5 U/mL Benzonase nuclease (Sigma-Aldrich, Saint Louis, MO, Catalog #E8263) overnight at 37 °C to digest free DNA. Phage was precipitated by adding 0.5 M NaCl + 4% w/v polyethylene glycol (PEG), molecular weight 8000 (Sigma-Aldrich, Saint Louis, MO, Catalog #PHR2894) overnight at 4 °C. Precipitated phage was then pelleted by centrifugation (14,000 x *g*, 20 min, 4 °C) and washed in 30 mL of Tris-EDTA buffer (10 mM Tris-HCl, 1 mM EDTA, pH 8.0). Then, the resuspended phage was re-pelleted by centrifugation (14,000 x *g*, 20 min, 4 °C) and resuspended in buffer appropriate to that phage/experiment and dialyzed against 4 L of the same buffer 3-4 times through a 10 kDa dialysis membrane to remove residual salts and PEG. This was repeated twice so that phages underwent two rounds of PEG precipitation (ThermoFisher Scientific, Waltham, MA, Product #A52972).

### Dynamic light scattering

All DLS size distributions were obtained using the Nano-Series Zeta Sizer (Nano-NS ZEN3600, Malvern Instruments, Worcestershire, United Kingdom) equipped with a 633 nm laser. Measurements were obtained at 25 °C at a backscattering angle of 173°. Each individual DLS measurement (single replicate) reported in this study is the average of 11 ten-second measurements obtained after a 1-minute vortexing period on half-speed, 1-minute equilibration period in the instrument, and with variable attenuation to generate count rates >100 kcps. Additional resuspension was performed in some cases due to sedimentation. Phage diffusion coefficients were calculated from auto-correlated light intensity data, and hydrodynamic diameters (D_H_) were calculated using ZetaSizer version 7 software with the Stokes-Einstein equation.

### Plaque assays

Plaque assays were used to quantify the number of infectious phage particles. We used a spot-dilution double-agar overlay technique. 100 μL of mid-log phase bacteria was added to 5 mL of top agar (5 g/L agar, 10 g/L tryptone, 10 g/L NaCl). Magnesium sulfate and calcium chloride were added to a final concentration of 20 mM. The mixture was poured onto non-selective LB-agar plates and allowed to solidify and dry briefly (5 min). Serial dilutions of phage were prepared in SM buffer (50 mM Tris-HCl, 100 mM NaCl, 8 mM MgSO_4_, 0.01% w/v gelatin, pH 7.5) and 10 μL of each dilution was spotted onto the top agar, incubated at 37 °C overnight, and plaques were counted.

### Transmission electron microscopy

The size and morphology of phages were examined with transmission electron microscopy (TEM) using a JEOL JEM1400 (JEOL USA Inc., Peabody, MA) at 80 kV. 5 μL of diluted phage solution was dropped onto carbon-coated copper grids (FCF-200-Cu, Electron Microscopy Sciences, Hatfield, PA). After 3 minutes, the grid was dipped into a ddH_2_O droplet. 1% uranyl acetate was dropped onto the sample for staining and allowed to dry for 15 minutes before performing microscopy.

### Sonication

500 μL of LPS5 at 10^10^ PFU/mL in SM buffer was sonicated using a probe sonicator (Branson Sonifier 250, Branson Ultrasonics, Danbury, Connecticut, USA) at maximum intensity for 10 seconds.

### Phage time series

1.5 mL of freshly-prepared CYPHY phages were prepared in SM buffer and dialyzed against 4 L of either SM buffer or PBS 4 times. Phages were then filtered through 0.22 um PES membranes, titered by plaque assay, and then diluted to a final concentration of 10^9^ PFU/mL. Phages were stored in 2 mL polypropylene tubes at 4 °C or 37 °C with weekly monitoring by triplicate DLS and 6 replicates of plaque assay. Belgian cohort phages were produced, filtered, stored for 1 month until all phages had been produced, and then monitored by triplicate DLS.

### Phage stability in different buffers

400 μL of CYPHY phage were aliquoted to 1.5 mL polypropylene tubes at a concentration of 1.25 x 10^8^ PFU/mL in various buffers as described in the text. Paired samples were incubated at 4 °C and 37 °C. Plaque assays and DLS size measurements were performed after 1 week of incubation to compare storage at 4°C vs. 37°C.

### Oxidation

Hydrogen peroxide was diluted from a 30% w/v stock into Tris-Mg buffer (50 mM Tris-HCl, 100 mM NaCl, 8 mM MgSO_4_, pH 7.5) and mixed with phage to a final concentration of 0.002% to 20% w/v peroxide. Peroxide-phage mixtures were mixed by pipetting. Reactions proceeded for 20 minutes before dialysis against Tris-Mg buffer in micro-dialysis cassettes (Xpress Microdialyzer MD300, 6-8 kDa, Scienova GmbH, Jena, Germany).

### *In vivo* murine full-thickness wound infection topical phage therapy model

All male C57BL/6J mice used for the *in vivo* wound infection phage therapy experiment were purchased from The Jackson Laboratory (Bar Harbor, ME). All experiments and animal use procedures were approved by the Institutional Animal Care and Use Committee (IACUC) at the School of Medicine at Stanford University. The study design was adapted from previously published work(37–39).

Briefly, 7–8-week-old male mice were anesthetized using 3% isoflurane. Mice dorsum were shaved using a hair clipper and depilated using Nair hair removal cream (Church and Dwight, Ewing, NJ). The shaved area was cleaned with betadine (Purdue Frederick Company, Catalog # 19-065534) and alcohol swabs (Coviden WEBCOL). Mice received 0.1-0.5 mg/kg slow-release buprenorphine (ZooPharm, Wedgewood Pharmacy, Swedesboro, NJ) subcutaneously as analgesic. Bilateral dorsal full-thickness wounds were created using 6-mm biopsy punches to outline the wounding area, and the epidermal and dermal layers (up to the fascia) were excised using sterilized scissors. The wounds were covered with Tegaderm (3M, Catalog # 1642W). Luminescent PAO1:lux was grown under antibiotic selection (100 μg/mL carbenicillin, 12.5 μg/mL kanamycin) in LB at 37 °C with shaking until mid-log phase. The inoculum was then diluted to 2.5 x 10^4^ CFU/mL in PBS and verified by plating. Mice received 10^3^ CFU/wound via injection onto each wound under the Tegaderm patch. Control mice were treated with SM buffer. Two hours post-infection, all mice received the first treatment. 24 hours after the first treatment, all mice received a second treatment dose. Mice were weighed daily and provided with Supplical Nutritional Supplement Gel (Henry Schein Animal Health Catalog # 0409-4888-10). Three days post-infection, mice were euthanized by CO_2_ chamber and cervical dislocation and the wound bed was excised, homogenized, and plated for CFUs onto LB agar plates.

### Statistical Analysis

All column graphs and statistical analyses were performed using Prism version 9 software (GraphPad, Boston, MA). Statistical significance was tested using two-tailed Pearson’s test for simple linear regressions, two-way ANOVA with Tukey correction for multiple groups, and one-way ANOVA with Tukey correction in the *in vivo* study. Mean ± SD was depicted to describe spread of data unless otherwise indicated.

DLS spectra of perturbed phage samples were compared to control phage samples by computing the sum of the absolute values of bin-wise differences between the two histograms on the size-intensity spectra in log scale (AUCΔ). Simple linear regression was used to develop standard curves. This analysis can be replicated for any two DLS spectra with matching titer measurements using the Phage-ELF tool available at https://jp22.shinyapps.io/shinyapp/ or R code available at https://github.com/jpourtois/phageELF.

## We gratefully acknowledge the following funding

National Institutes of Health grant R01 HL148184-01 (PLB)

National Institutes of Health grant R01 AI12492093 (PLB)

National Institutes of Health grant R01 DC019965 (PLB)

Cystic Fibrosis Foundation grant (PLB)

Emerson Collective grant (PLB)

Stanford University Medical Scientist Training Program grant T32-GM007365 (TD)

Stanford Interdisciplinary Graduate Fellowship, Gold Family Graduate Fellow (TD)

## Data availability

The datasets generated during and/or analyzed during the current study are available from the corresponding author on reasonable request.

## Material availability

All unique biological materials (phages) are readily available from the authors, except in the case of some Belgian phages, which have decayed.

## Code availability

The Phage-ELF software is available as a web application at https://jp22.shinyapps.io/shinyapp/. The R code is available for download at https://github.com/jpourtois/phageELF.

