## Supplemental Figures and Tables for "Rapid assessment of changes in phage bioactivity using dynamic light scattering"

**This PDF file includes:**

Figures S1 to S2
Tables S1 to S2

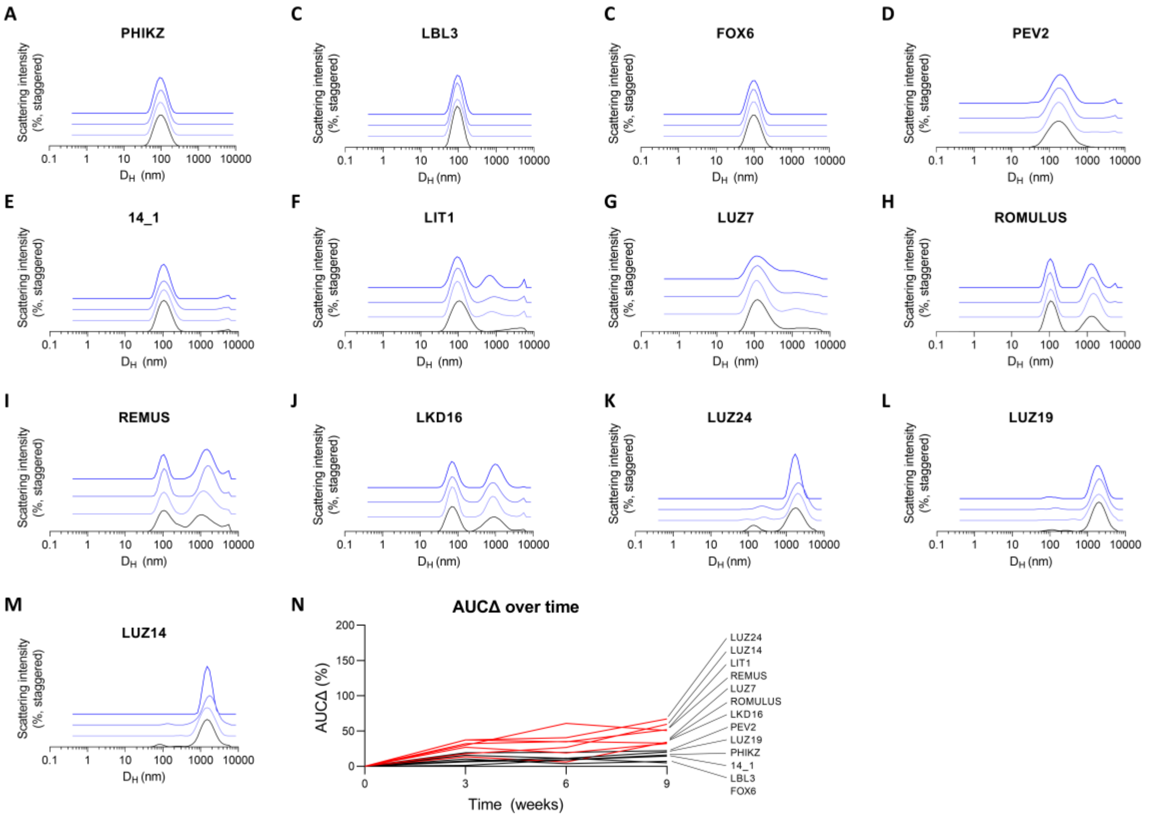


**Fig. S1. Phages spontaneously fragment and aggregate in refrigerated storage over time (Belgian cohort).**

(**A** through **M**) DLS spectra of phages from the Belgian cohort (BC) over a monitoring period of two months. Shown are averages of n=3 DLS measurements per phage. n=13 phages were assessed. Darker colors and vertical staggering are used to show progression in time. Few phages remained largely intact (**A** through **C**). Most phages aggregated over the monitoring period (**D** through **M**). (**N**) AUCΔ identifies the most-changed phage over the monitoring period and is consistent with our qualitative assessment of the DLS spectra.

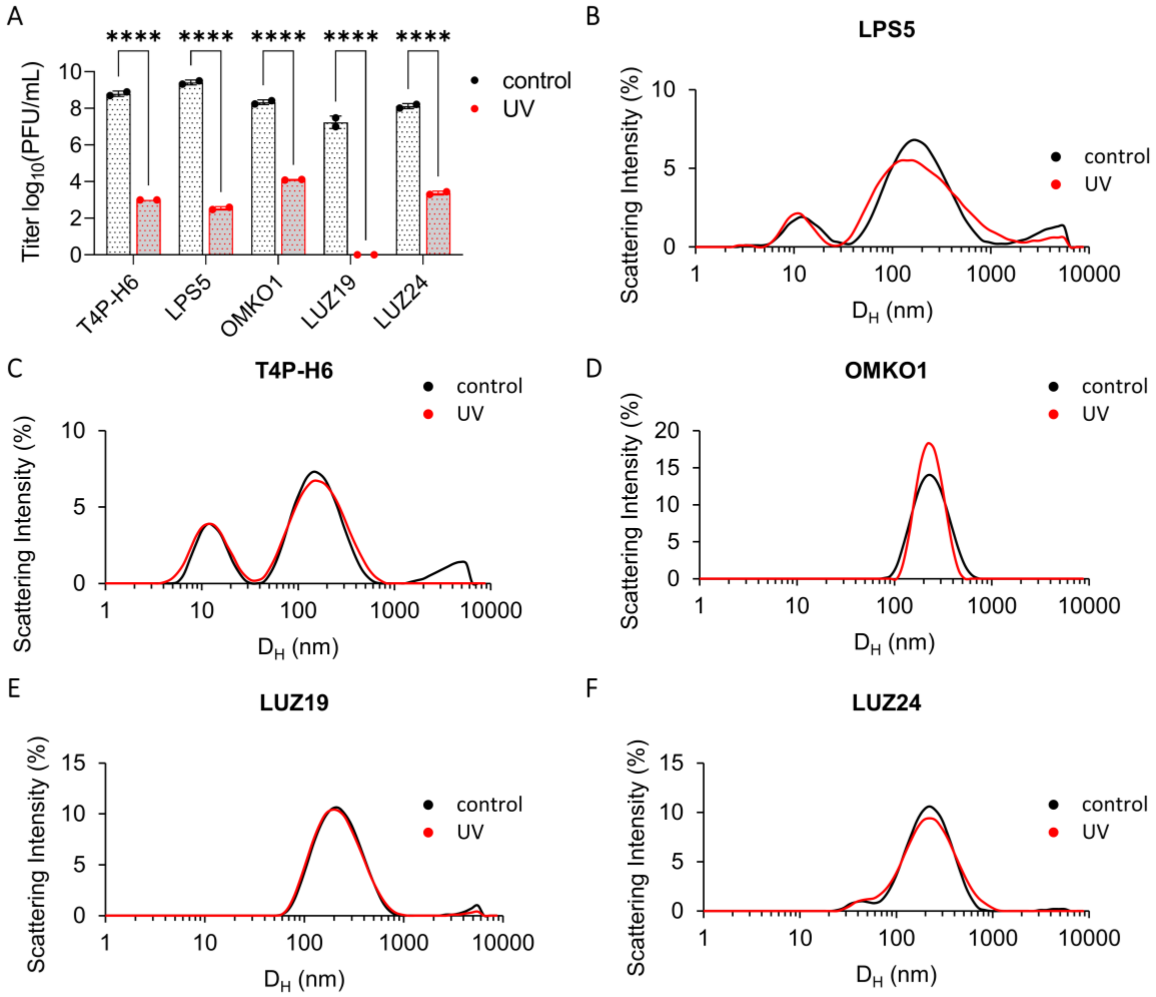


**Fig. S2: DLS captures changes in phage size, but it does not capture genomic damage.**
(**A**) Titer of phages after irradiation for 20 minutes with germicidal UV-C light. Results are from one experiment. Titer was measured with n=2 plaque assays per phage per condition. Two-way ANOVA with Tukey correction. **** = $p < 0.0001$. (**B** through **F**) DLS spectra of phages before and after irradiation. Shown are averages of n=3 DLS measurements per phage per condition.

**Table S1. Physical and biological characteristics of CYPHY phages.**

| Phage | T4P-H6 | LPS5 | OMKO1 |
|---|---|---|---|
| **Plaque characteristics** | Clear, well-circumscribed<br>Size: 0.3 – 1 mm | Clear, well-circumscribed<br>Size: 1.5 – 2.0 mm | Clear, well-circumscribed<br>Size: 0.5 – 1.5 mm |
| **Dimensions** | Tail length: 190 ± 5 nm<br>Tail width: 16 ± 2 nm<br>Head length: 74 ± 2 nm<br>Head shape: isometric | Tail length: 140 ± 2 nm<br>Tail width: 20 ± 1 nm<br>Head length: 70 ± 3 nm<br>Head shape: isometric | Tail length: 212 ± 2 nm<br>Tail width: 28 ± 1 nm<br>Head length: 134 ± 4 nm<br>Head shape: isometric |
| **Morphology** | *Siphoviridae* | *Myoviridae* | *Myoviridae* |

**Table S2. Phages used in this study.**

| Phage | Taxonomy | Bacterial Host | Morphology | genome size (kb) | dim. (nm) capsid / tail |
|---|---|---|---|---|---|
| LUZ14 | *Autographiviridae* | *P. aeruginosa* C1 | podovirus | ~43 | 62/12 |
| LUZ19 | *Autographiviridae, Phikmvvirus* | *P. aeruginosa* PA01 K | podovirus | 43.5 | 65/12 |
| LUZ24 | *Bruynoghevirus* | *P. aeruginosa* Li010 | podovirus | 45.6 | 63/12 |
| LKD16 | *Autographiviridae, Phikmvvirus* | *P. aeruginosa* GHB15 | podovirus | 43.2 | 65/12 |
| LUZ7 | *Schitoviridae, Luzseptimavirus* | *P. aeruginosa* Br257 | podovirus | 74.9 | 76/30 |
| PEV2 | *Schitoviridae, Litunavirus* | *P. aeruginosa* PA01 K | podovirus | 72.7 | 70/30 |
| LIT1 | *Schitoviridae, Litunavirus* | *P. aeruginosa* US449 | myovirus | 72.5 | 74/30 |
| LBL3 | *Pbunavirus* | *P. aeruginosa* C1 | myovirus | 64.4 | 73/148 |
| 14_1 | *Pbunavirus* | *P. aeruginosa* Li010 | myovirus | 66.2 | 73/148 |
| PhiKZ | *Phikzvirus* | *P. aeruginosa* Aa245 | myovirus | 280.3 | 145/200 |
| Romulus | *Herelleviridae, Silviavirus* | *S. aureus* (broad host range) | myovirus | 131.3 | 90/204 |
| Remus | *Herelleviridae, Silviavirus* | *S. aureus* (broad host range) | myovirus | 134.6 | 90/204 |
| Fox6 | *Carmasinavirus* | *X. campestris* pv. *campestris* I11008 | myovirus | 61.1 | 78/156 |
| OMKO1 | *Phikzvirus* | *P. aeruginosa* PAO1 | myovirus | 281.8 | 134/212 |
| LPS5 | *Pakpunavirus* | *P. aeruginosa* PAO1 | myovirus | 93.1 | 70/140 |
| T4P-H6 | *Nipunavirus* | *P. aeruginosa* PA14 | siphovirus | 57.4 | 74/190 |
| T2 | *Straboviridae, Tequatrovirus* | *E. coli* B | myovirus | 163.8 | 111/78 |
| T3 | *Autographiviridae, Teetrevirus* | *E. coli* B | podovirus | 38.3 | 60/30 |
| T4 | *Straboviridae, Tequatrovirus* | *E. coli* B | myovirus | 168.9 | 111/78 |
| T6 | *Straboviridae, Tequatrovirus* | *E. coli* B | myovirus | 170 | 120/86 |
| T7 | *Autographiviridae, Tespetimavirus* | *E. coli* B | podovirus | 39.9 | 55/29 |
